# Elastin Recoil is Driven by the Hydrophobic Effect

**DOI:** 10.1101/2023.03.07.531587

**Authors:** Nour M. Jamhawi, Ronald L. Koder, Richard J. Wittebort

## Abstract

Elastin is an extracellular matrix material found in all vertebrates. Its reversible elasticity, robustness and low stiffness are essential for the function of arteries, lungs, and skin. It is among the most resilient elastic materials known: During a human lifetime, arterial elastin undergoes in excess of 2x10^9^ stretching/contracting cycles without replacement and slow oxidative hardening has been identified as a limiting factor on human lifespan. For over fifty years, the mechanism of entropic recoil has been controversial. Herein, we report a combined NMR and thermomechanical study that establishes the hydrophobic effect as the primary driver of elastin function. Water ordering at the solvent:protein interface was observed as a function of stretch using double quantum ^2^H NMR and the most extensive thermodynamic analysis performed to date was obtained by measuring elastin length and volume as a function of force and temperature in normal water, heavy water and with co-solvents. When stretched, elastin’s heat capacity increases, water is ordered proportional to the degree of stretching, the internal energy decreases, and heat is released in excess of the work performed. These properties show that recoil in elastin under physiological conditions is primarily driven by the hydrophobic effect rather than by configurational entropy as is the case for rubber. Consistent with this conclusion are decreases in the thermodynamic signatures when co-solvents that alter the hydrophobic effect are introduced. We propose that hydrophobic effect-driven recoil, as opposed to a configurational entropy mechanism, where hardening from crystallization can occur, is the origin of elastin’s unusual resilience.

**Significance:** Elastin, found in tissues that require reversible elasticity, has low stiffness and great resiliency. It is a self-assembled material that has been a target for regenerative medicine. However, the basis for its elasticity has been controversial for more than 50 years. Formed from a hydrophobic protein with an equivalent mass of water, the controversy is whether recoil is driven by entropy gain of the protein and/or the water. We demonstrate that matrix water is progressively ordered upon stretching and that the thermodynamics of elastin recoil are those of the hydrophobic effect and different from those of rubber. We conclude that recoil is primarily driven by the hydrophobic effect and suggest that this accounts for elastin’s low stiffness and high resilience.

## Introduction

Although elastin has been identified as one of the limiting factors on human lifespan (1), its molecular mechanism of entropic recoil is not well-understood and has been a source of controversy for over fifty years (2-12). A widely accepted hypothesis is that elastin is like rubber and an increase in the polymer’s configurational entropy drives recoil (2, 3, 13). Unlike rubber, elastin gains elasticity only when swollen with an approximately equal mass of water (14) and its physiological function requires different mechanical properties: First, elastin recoil is fully reversible on the timescale of a heartbeat. This enables the elastic energy stored in arteries when the heart contracts to be fully recovered when the heart relaxes between beats. Second, elastin is highly compliant, with a Young’s modulus (∼ 0.4 MPa) that is an order of magnitude less than a typical rubber (∼ 10 MPa) (15). Third, elastin is far more robust than rubber (16), undergoing more than 2x10^9^ expansion/contraction cycles without appreciably lengthening during a normal human lifespan and failure of any of these properties results in hypertension which in turn causes vascular calcification, ventricular hypertrophy, renal dysfunction and stroke.

The elastic matrix is formed when tropoelastin, one of the most hydrophobic proteins found in nature, is exported to the extracellular matrix and undergoes a liquid-liquid phase separation followed by enzymatic cross-linking (17, 18). Tropoelastin is organized in alternating proline-rich hydrophobic domains and alanine-rich crosslinking domains. (Figure 1a) (19). In 1970, Weis-Fogh (20) found that the amount of heat released when elastin is stretched is in large excess over the applied work (*ΔG*) and concluded that, unlike rubber (21, 22), elastin’s internal energy decreases when stretched (4, 9, 23). It was hypothesized that the solvent exposed surface area of hydrophobic droplets increases when deformed and ordering of water at the hydrophobic surface of elastin droplets is increased, i.e., spontaneous recoil is driven by the hydrophobic effect. Flory and Hoeve rejected the droplet model (2) in favor of a rubber-like network and pointed out that the thermodynamic analysis of Weis-Fogh (4) had neglected the contribution to the change in internal energy from a volume change that could occur when the sample is stretched, and they found that elastin’s volume and internal energy are constant when stretched in aqueous glycol (3:7) or 30% PEG. They concluded that elastin is like rubber, i.e., configurational entropy of the polymer drives recoil (2, 3). However, their analysis neglected the effect of glycol or PEG on water:protein interactions (24, 25) and it was subsequently reported (9) that elastin’s volume is constant when stretched in water.

**Figure 1.**
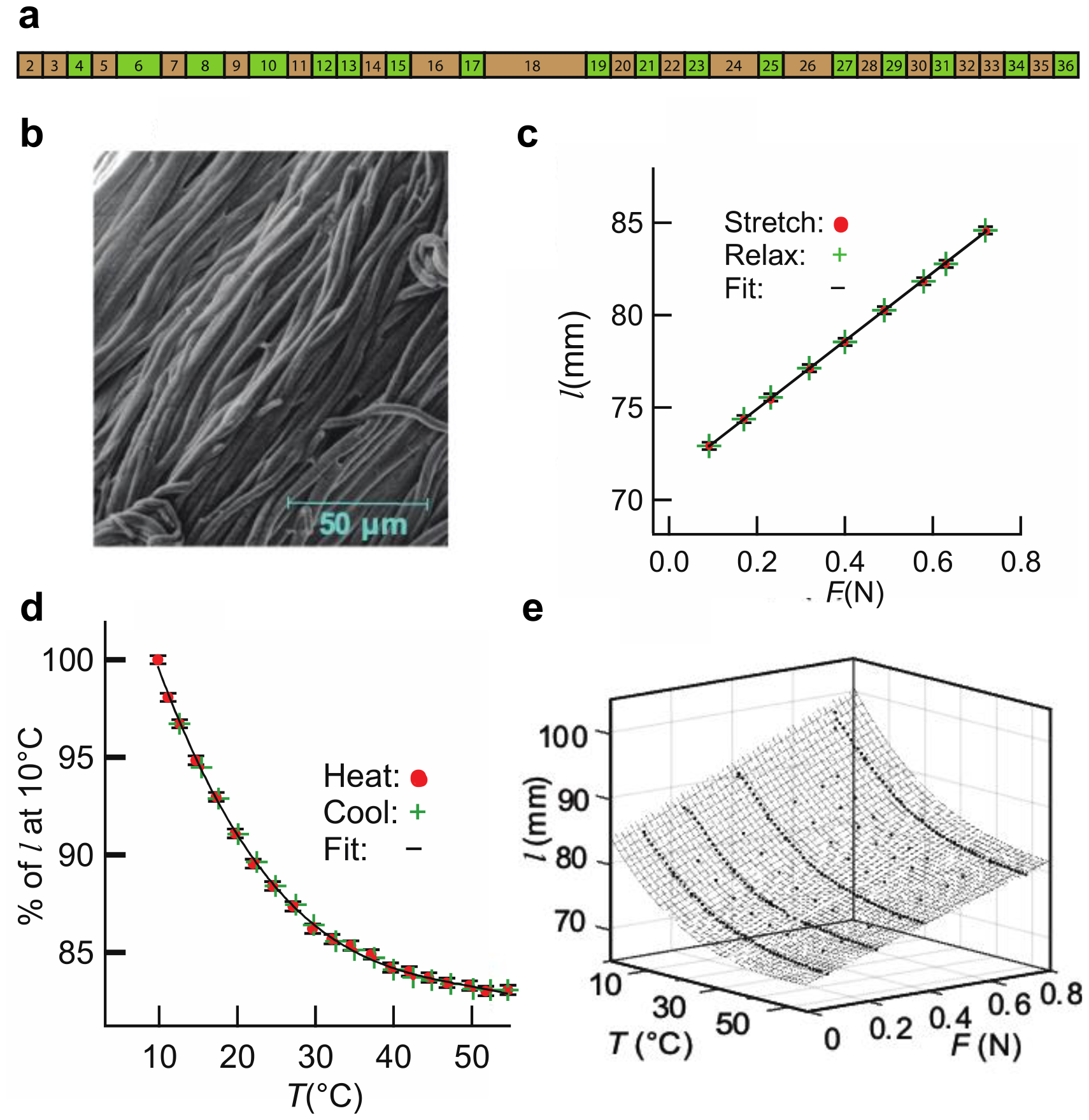
(**a**) Domain architecture of bovine elastin. Brown rectangles represent hydrophobic domains and green rectangles represent crosslinking domains. (**b**) SEM of purified bovine elastin showing the uniform fibrous structure found in natural elastin. (**c**) Reversibility of the fiber length from two stretch and contract cycles. The fit to two cycles (line) is linear. (**d**) Reversibility from heating and cooling. The fit (line) is a polynomial 3^rd^-order in *T*. (**e**) Surface plot, *l(F,T)*, obtained by combining data from multiple *l* vs. *F* and *l* vs. *T* experiments. The fit (lines) is a polynomial 1^st^-order in *F* and 3^rd^-order in *T*.

In this report, stretch-induced ordering of water (^2^H_2_O) in the elastin matrix was quantified using double quantum (2Q) filtered ^2^H NMR, (26, 27) and a complete thermodynamic study of elastin’s mechanical properties was obtained. The NMR experiment uses the property that isotropic molecular tumbling in bulk water (^2^H_2_O) averages the ^2^H quadrupole coupling to zero whereas tumbling of water at a hydrophobic surface is anisotropic and the coupling is non-zero. Using the methods of multiple quantum spectroscopy, the bulk water signal is eliminated from detection and only the 2Q signal from ordered water is observed as the sample is stretched (28). We have determined elastin’s thermodynamic properties with a custom-made thermomechanical apparatus. The length and volume of a purified elastin sample were measured with a high-resolution camera as a function of applied force and temperature (from 3°C to 55°C) in several aqueous solvents from which virial equations of state were constructed and the relevant thermodynamic properties, *ΔG, ΔS, ΔH, ΔU* and *ΔC*_*p*_, were calculated as functions of temperature and strain. Of particular interest are the dependencies of the heat capacity, enthalpy and entropy with temperature and strain. When water molecules contact a hydrophobic surface, they are weakly ordered (29), the H-bond angle becomes more ideal and the H-bond length decreases (30). Consequently, entropy is lower and heat is released. As the temperature is increased, water is less ordered and the H-bond lengthens, i.e., *ΔS* and *ΔH* both increase (become less negative) resulting in a comparatively small change in *ΔG* (2*9)*, known as entropy-enthalpy compensation. With reduced heat loss, the heat capacity increases (31).

## Results

### Reversibility and the equation of state

Figure 1a depicts the domain structure of bovine elastin (747 residues) and Figure 1b depicts a scanning electron micrograph demonstrating that the fibrous structure of natural elastin has been preserved in the purification procedure used here. (32) A requirement in our thermodynamic analysis is reversibility of the length following changes in the applied force or temperature. Figures 1c and 1d show that elastin’s length subjected to stretch/contract and heat/cool cycling is, as expected, fully reversible on the time scales of ∼5 minutes for length changes and 3-5 minutes for temperature changes. From 3°C to 55°C, the decrease in length, *l*, is non-linear and reaches a temperature-independent length above ∼50 °C (Figure 1d). The variation of *l* with force, *F*, is linear as previously reported. (9) The *l(F,T)* surface (Figure 1e), was obtained by combining multiple *l(F)* and *l(T)* data sets and fitting 308 data points (elastin in water) to a virial equation of state (equation 8). The RMSD of 0.26 mm is comparable to the estimated accuracy of the length measurements. Fit parameters for elastin in water and in other solvents are listed in Table S1. The equation of state (equation 8) is conveniently expressed in the form of Hooke’s law (equation 11) with relaxed length, *l*_*rlx*_ (equation 9) and force constant, *k*_H_ (equation 10).

### Elastin’s mechanical and thermodynamic properties in pure water and comparison to natural rubber

The thermo-mechanical parameters obtained from purified elastin in water at 10°C, 35°C and 45°C are summarized in Figure 2. Solid lines were calculated using equations 9-17 and the virial equation coefficients in Table S1. Error bars, shown at regular intervals along the solid lines, are standard deviations of distributions of the thermodynamic property obtained by propagating uncertainties in the virial coefficients into standard deviations of each property (33) as described in the **Materials and Methods** and used throughout this manuscript. So that data from different experiments could be directly compared, a single sample with randomly oriented fibers was used for all experiments and its integrity was confirmed by the reproducibility of the length versus temperature curves obtained following experiments in different solvents, Figure S6. Samples with different fiber orientations potentially yield somewhat different results, however, the Young’s modulus, 0.4 MPa at 37°C (Figure 2a), is in good agreement with the value of 0.45 MPa previously determined from unpurified, single-fiber samples (23). As temperature is raised, the change in *E* results from a force constant that increases linearly and a relaxed length that decreases in a non-linear way (Figure 2b) indicating increased attractive forces in the elastin matrix. The relaxed length reaches a constant value at temperatures above ∼50°C (Figure 2b).

**Figure 2:**
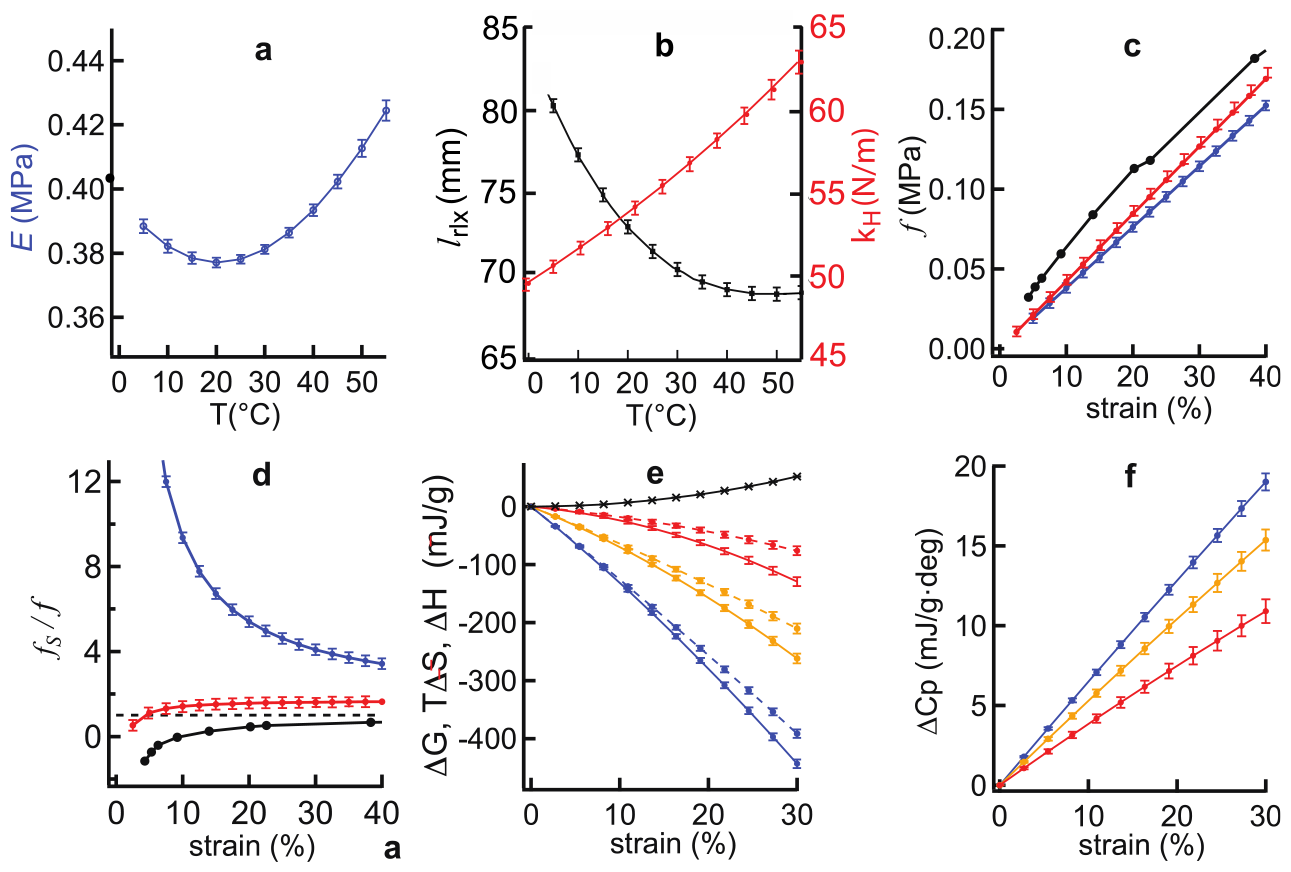
Temperature dependence of elastin’s mechanical and thermodynamic properties. (**a**) Young’s modulus, (**b**) the relaxed length, *l*_rlx_ (▀), and the force constant, k_H_ 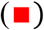. (**c**) Tension, *f*, as a function of strain for elastin at 30°C 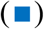, elastin at 50°C 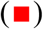 and rubber at 30° (▀). (**d**) Ratio of the entropic tension to the total tension, *f*_S_/*f*, as a function of strain for elastin at 30°C 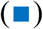, elastin at 50°C 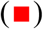 and rubber at 30°C (▀). (**e**) Elastin *ΔG* (▀), *ΔH* (⋯) and *TΔS* (−) vs strain at 10°C 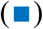, 25°C 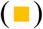 and 45°C 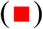. *ΔG* values at 10°C, 25°C and 45°C overlap. **(f**) Elastin ΔC_*p*_ vs strain at 10°C 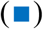, 25°C 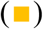 and 45°C 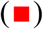.

Elastin and rubber tension (force per unit area) measured at constant *T* and *P* can be directly compared. Tension, *f*, in elastin is less than that of rubber and varies linearly with strain (Figure 2c). With equation 7, *f* can be separated into entropic and enthalpic components (*f = f*_S_ *+ f*_H_) (13, 23). Ratios of *f* and *f*_S_ were calculated for elastin using equation S4 with the virial coefficients listed in Table S1 for elastin and values of *f* and *f*_S_ from the data of Shen, *et*.*al*. (21) for rubber. The ratios of *f*_S_ /*f* for elastin at 30°C and 50°C are depicted as a functions of strain and compared to rubber at 30°C (Figure 2d) (21). The horizontal line, *f*_S_/*f* = 1, represents an ideal elastomer in which case the force is entirely entropic. At 30°C and strains greater than 20%, rubber is approximately ideal (*f*_*S*_*/f* ∼0.8) whereas elastin is far from ideal at all strains (*f*_*S*_*/f* > 4). When the temperature of elastin is raised to 50°C, the entropic contribution to the total tension approaches that observed in rubber. Because the entropic tension is large compared to the net tension at physiological temperatures, the entropic and enthalpic tensions are opposed in elastin and the magnitude of the enthalpic tension, *f - f*_S_, is almost as large as the entropic component and decreases in parallel with *f*_S_ as the temperature is increased.

Elastin’s thermodynamic properties as functions of strain and temperature are depicted in Figures 2e and 2f. The expressions used to obtain *ΔG, ΔH, ΔS* and *ΔCp* from the virial equation of state, are derived in the materials and methods section and apply to an open system at constant temperature and pressure that is in equilibrium with the surrounding water. The thermodynamic parameters determined in this way are for the entire system and change as the temperature and sample length are varied. Importantly, the heat to work ratios, *ΔH/ΔG*, determined at 23°C using our method when elastin is stretched to strains of 10%, 20% and 30% are -23±0.3, -11.9±0.3, and -8.1±0.2, respectively, which are in agreement with values previously determined calorimetrically (9).

Elastin’s thermodynamic properties are fully consistent with the conclusion that stretch increases the exposure of elastin’s predominantly hydrophobic residues to water and this contributes to the net thermodynamic force that drives recoil: when elastin is stretched from the relaxed state, *ΔH* and *TΔS* are negative at all values of strain and temperature and their small net difference, *ΔG*, gives elastin its physiologically important property of high compliance (low Young’s modulus). The magnitudes of *ΔH* and *TΔS* decrease substantially as the temperature is raised while *ΔG* is nearly independent of temperature. At temperatures below 45°C, spontaneous recoil in elastin is endothermic and entropy driven with a free energy change that is less than the heat liberated. Elastin’s heat capacity (Figure 2f) has not been previously determined and we find that *ΔC*p increases when strain is applied. The pattern of changes in *ΔG, ΔH* and *TΔS* (Figure 2e) and the increase in heat capacity (Figure 2d) are all characteristic features of the transfer of hydrophobic substances into an aqueous environment (29, 30, 34). The decrease in the amount of ordered water, and its concomitant decrease in volume, are expected consequences of the hydrophobic effect.

### Volume expansion and water uptake is small when elastin is stretched

Because heat is in large excess of the work performed when stretched, it was concluded that, unlike rubber, elastin’s internal energy decreases when stretched (4, 9). McCrum *et al*., in a series of papers, pointed out that this conclusion neglected a change in volume that could occur when elastin is stretched and proposed that the release of heat from water uptake associated with an increased volume could account for the large release of heat when elastin is stretched (2, 5-7). However, it was subsequently reported that the increase in elastin’s volume when extended by 20% was negligible (9).

To resolve this disagreement, we measured elastin’s volume as a function of strain and temperature, Figure 3a, and applied the thermodynamic relation *ΔU = ΔH* - *PΔV*. Volume was estimated from the digital images of the sample. The coefficients of expansion calculated from the volume data, 14.8x10^−3^ at 15°C and 3.4x10^−3^ at 38°C, are in good agreement with previous determinations (10). At temperatures from 10°C to 45°C, elastin’s volume expands by 5% to 6% when stretched isothermally to a strain of 15%. At 37°C, the change in internal energy from expansion (-15±2 mJ/g) is small compared to that from enthalpy (-144.5±6.3 mJ/g) and the reduction in internal energy (-159.5± mJ/g) is, in fact, large compared to the stretch work (35.5±0.3 mJ/g). As temperature is increased, heat loss is reduced and at 50°C, the magnitudes of the enthalpy change, and work are comparable (Table 1). Like the effect of temperature, the internal energy loss is also reduced when co-solvents, 0.3 m sodium sulfate or 30% PEG, are added (Figure 3c).

**Table 1:**
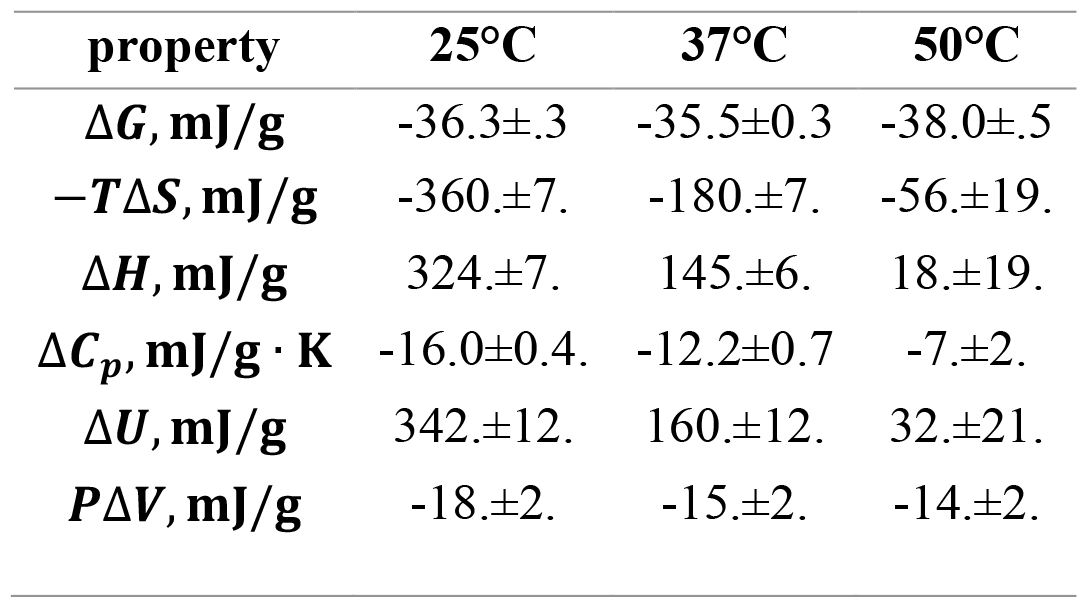
Thermodynamic properties of elastin recoil from 25% strain in water at three temperatures. Sample strain was 25%.

**Figure 3:**
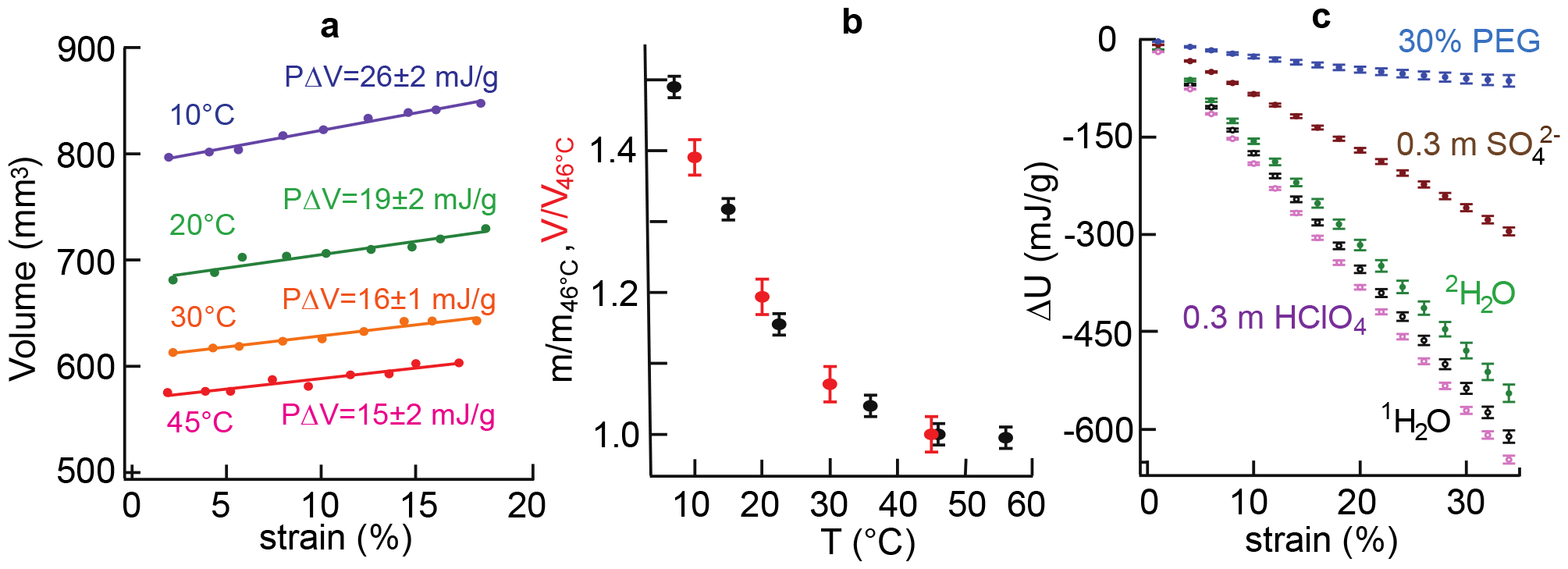
(**a**) Elastin volume as a function of strain and temperature. Lines are linear fits to the volume vs strain data and the indicated values of *PΔV* are per gram of protein dry weight at 15% strain. (**b**) Temperature dependence of the mass (•) and volume 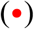 of elastin samples normalized to their values at 46°C. (**c**) Strain induced values of Δ*U* for elastin at 20°C in the indicated solvents.

Elastin’s volume increases by a large amount, 40%, when cooled (Figure 3a) and we find that the mass and volume of an elastin sample vary in direct proportion with one another (Figure 3b). We conclude that the increase in volume when elastin is stretched or cooled is due entirely to the influx of water, i.e., the changes in the equilibrium volume, *dV*_*eq*_, and the number of moles of water of hydration, 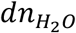, are proportional and related by, 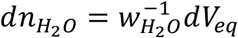, where 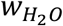 is the partial molar volume of water in the elastin matrix. With the known dry mass of the vacuum-dried sample, we find that elastin is 48% and 63% water by weight at 45°C and 10°C, respectively.

### Co-solvents and solvent deuteration alter elastin’s mechanical and thermodynamic properties

To further confirm the conclusion that the hydrophobic effect is the primary driver of elastic recoil, we have studied elastin’s thermo-mechanics with three cosolvents, Na_2_SO_4_, 20kD PEG and NaClO_4_, and when ^1^H_2_O is replaced by ^2^H_2_O, each of which changes water and/or the protein in different ways (Figure 4). The anions, SO_4_^2-^ and ClO_4_^-^, are strong Hoffmeister ions that are, respectively, kosmotropic and chaotropic, and their effects dominate those from the counterion, Na^+^, a weak kosmotrope.

**Figure 4:**
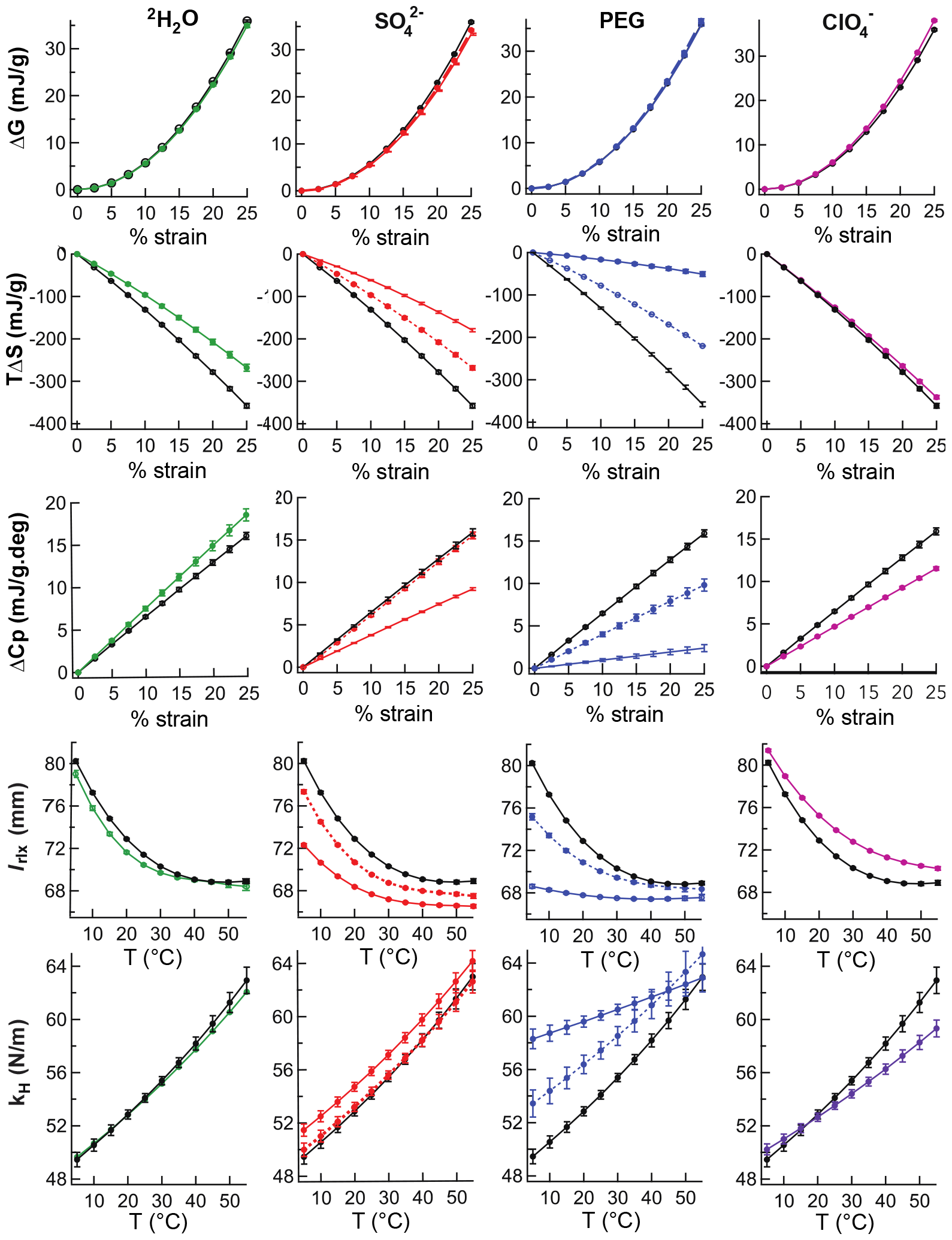
Thermomechanical properties of elastin in different aqueous solutions. Columns in the figure are the different solvents: ^2^H_2_O (−), Na_2_SO (⋯ 0.1 m and − 0.3 m), 20 kDa PEG (⋯ 15% and − 30%) and NaClO_4_ (−0.3 m). Rows in the figure are thermodynamic parameters (*Δ*G, *TΔ*S and *ΔC*_*p*_) are plotted versus % strain at 25°C and the mechanical parameters (*l*_*rlx*_ and *k*_H_) are plotted versus temperature. For reference, the data from elastin in pure water (−) is shown in each graph.

PEG and SO_4_^2-^ are widely used to prepare crystalline proteins in their native forms (35). They alter the water:protein interface (36) and increase attractive forces in proteins because they are preferentially excluded from the protein’s surface (24, 25, 37). Sulfate increases water’s surface tension. This raises the energy required to make the cavity required to accommodate exposed hydrophobic side chains causing the protein to bury hydrophobic residues and become more compact (38). High molecular weight PEG is preferentially excluded from the protein surface because its large size limits penetration into the volume surrounding a protein, known as the exclusion (24, 25) or depletion zone (39). As shown schematically in references 39 (Figure 3) and 41 (Figure 6), when a protein associates or compacts, the exclusion volume is reduced and the volume available to PEG increases, giving it reduced free energy and greater entropy (39). We find that when Na_2_SO_4_ or PEG is added at 25°C, all changes in the thermomechanical properties, Figure 4, are similar to those found in elastin in water when the temperatures is raised above 45°C, Figures 2e and 2f. The relaxed length and volume are reduced, the force constant is increased, the entropic force is decreased, the release of heat decreases, the increase in heat capacity is diminished, and the net driving force is nearly unchanged. For example, when elastin is stretched by 25% at 25°C in 30% PEG, the heat released decreases from 375 mJ/g to 10 mJ/g, Figure S2, the increase in *ΔC*_*p*_ is reduced from 17 mJ/K to 2 mJ/K, Figure 4, and the decrease in internal energy equals the stretch work.

Compared to sulfate, elastin is substantially less affected by 0.3 m ClO_4_^−^, a chaotrope, and changes in *l*_rlx_ and k_H_ are in the opposite direction, Figure 4. Perchlorate binds weakly to the backbone and denatures proteins. When negative charge is added to the nearly neutral elastin matrix, repulsive forces are introduced causing *l*_*rlx*_ to increase and *k*_H_ to decrease. This effect is small because elastin lacks significant 2° structure (40-43) – weak perchlorate binding to the backbone cannot significantly disrupt a disordered protein. Since 0.3 m ClO_4_^-^ has little effect on surface tension, the large and opposite effects from attractive forces seen with added SO_4_^2-^ are absent. However, when the concentration is increased to 1 m, perchlorate becomes kosmotropic with increased surface tension (44) and the changes in *l*_rlx_ and k_H_ seen at 0.3 m are reversed (Figure S1).

When stretched in ^2^H_2_O, *k*_H_ and *l*_rlx_ are smaller than in ^1^H_2_O, *TΔS* is less negative, *ΔC*_p_ rises faster and *ΔG* is unchanged (Figure 4). Similarly, when folded proteins are denatured in ^2^H_2_O rather than ^1^H_2_O, an increase in *ΔC*_p_ with constant *Δ*G (entropy-enthalpy compensation) is usually observed and this has been associated with an increase in solvent exposed, non-polar surface area (45, 46). These results are fully consistent with the greater solubility of non-polar molecules in ^2^H_2_O (47) and show that small changes in the strength of the hydrophobic effect have observable effects on the thermodynamics of elastin stretch and recoil. These surface effects are more important than, for example, stronger H-bonds in the protein which should have the opposite effect on the stretch *ΔH*. This is consistent with our NMR studies indicating a near absence of stable 2° structure in elastin (40, 48) and molecular dynamics simulations of an aggregated elastin-like peptide (11) indicating that putative β−turns stabilized by H-bonds have sub-nanosecond lifetimes. Each of these cosolvents alters the thermodynamics of stretching in elastin in a manner consistent with recoil primarily driven by the hydrophobic effect.

### Water is ordered when elastin is stretched

The thermodynamic experiments described above provide strong thermodynamic evidence for the primacy of the hydrophobic effect in elastin recoil. We next set out to directly observe water ordering – the hallmark of hydrophobic effect - upon stretching of elastin (29). To determine if water ordering in elastin varies with strain, we used double-quantum (2Q) filtered ^2^H NMR of elastin in ^2^H_2_O. In this experiment, the signal from bulk water which reorients isotropically is removed and only the 2Q signal from ordered water which reorients anisotropiocally and has a non-zero quadrupole coupling is detected (26, 27). The experiment is quantitated in terms of the water fraction, f_o_, that is ordered. Note that elastin’s mechanical properties in ^2^H_2_O are almost the same as in ^1^H_2_O (Figure 4). We find that the ordered water fraction, 2% in relaxed elastin, increases substantially to 81% when the sample is stretched by 27% (Figure 5a). This large increase is direct observation of the hydrophobic effect at work during elastin function.

**Figure 5:**
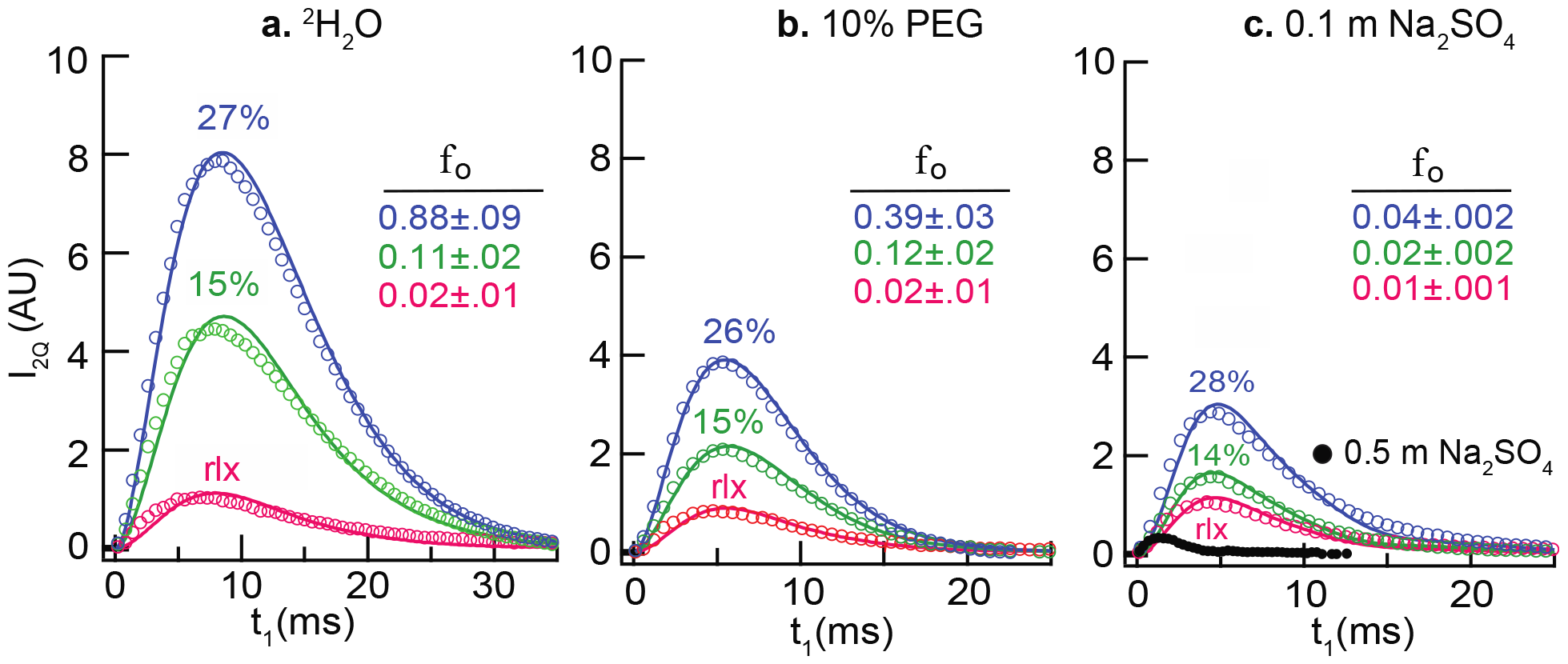
Plots of the deuterium 2Q NMR signal intensity, I_2Q_, as a function of the 2Q preparation time, t_1_. Open circles are the intensities of the 2Q filtered NMR signal with only every third data point shown. The ordered fraction, f_o_, was determined by least squares fitting (solid lines) to equation 18. Elastin fibers were hydrated in (**a**) ^2^H_2_O, (**b**) 10% PEG (w/w), (**c**) 0.1 m Na_2_SO_4_ and 0.5 m Na_2_SO_4_ at 28% stretch (black dots). The fibers are relaxed 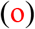, 14-15% stretched 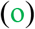, and 26-28% stretched 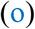.

Next, we observed the effects of cosolvent on water ordering: in 10% PEG, the ordered fraction is reduced ∼2-fold compared to pure ^2^H_2_O at each stretch (Figure 5b) and further lowered in 0.1m sodium sulfate (Figure 5c). In 0.5m sodium sulfate, ordered water is almost eliminated from observation in this experiment (Figure 5c). Overall, these results show that the amount of ordered water decreases with the addition of cosolvents that reduce solvent exposed surface area: Na_2_SO_4_ and PEG.

## Discussion

To understand the molecular basis of recoil in elastin, we have combined a complete study of elastin’s thermodynamics and measured water ordering in the elastin matrix at the molecular level. Under physiological conditions, elastin is a non-ideal elastomer, as summarized in Table 1. At 37°C, the entropic force that drives recoil, −*TΔS*, is five-fold greater than the magnitude of the net driving force, *ΔG*. In rubber, the entropic and the net driving force are approximately equal, as is the case for an ideal elastomer (22). Because −*TΔS* and the enthalpy change, *ΔH*, are opposed (opposite sign), elastin has high compliance. Otherwise, a heart with greater contractive force would be required to expand arteries. From Le Chatelier’s principle, the large heat absorption with recoil follows directly from the large decrease in the relaxed length when the temperature is raised (Figure 1d). At 50°C, the relaxed length reaches a constant value. Consequently, *ΔH* and *TΔS* for recoil from 25% strain are drastically reduced to 18.1±19 mJ/g and 56.3±19 mJ/g, respectively, from their values at 5°C, 760±19 mJ/g and 801±19 mJ/g. Since *ΔG* is nearly constant, -38.0±0.5 mJ/g at 50°C and -41.5±0.5 mJ/g at 5°C, there is significant enthalpy-entropy compensation. This and the stretch induced increase in the heat capacity, also observed here, are signature features of the hydrophobic effect.

With thermo-mechanics, we have confirmed that the heat released from stretch is in large excess over the total work performed, previously determined using a calorimeter (4, 9). In addition, we have determined that this process is fully reversible (Figure 1c). Thus, objections to this finding based on a time-scale argument (5) can be disregarded. Because *ΔH* from stretch is large and negative, Weis-Fogh concluded that, unlike rubber (22), elastin’s internal energy had decreased. Flory, however, observed no change in the volume or internal energy when elastin was stretched in 30% PEG and concluded that elastin is like rubber (2). While Weis-Fogh had assumed that volume was constant when stretched, Flory had assumed that PEG has negligible effect on water:protein interactions. To resolve this, we have measured elastin’s volume as a function of temperature and strain and used the general relation, Δ*U*= Δ*H* − *P*Δ*V*, to determine *ΔU* in pure water and in 30% PEG. We concur that *ΔV* in 30% PEG is negligible, however, *ΔV* is small in water and the contribution to the internal energy is less than 10% of *ΔH* (Figures 2 and 3). Thus, in physiological conditions, *ΔU* decreases when elastin is stretched. The difference between elastin in water and 30% PEG is primarily the difference in *ΔH* and not *ΔV*.

To better understand the effects of co-solvents, we have determined elastin’s thermo-mechanical properties and the ordered water fraction, f_o_, with three co-solvents. Sodium sulfate and PEG change the water:protein interface, do not bind to the protein and promote protein compaction while perchlorate binds to the protein, has little effect on water properties and leads to protein expansion. These expectations are fully borne out in changes to elastin’s thermo-mechanical properties (Figure 4), and to changes in the ordered water fraction, f_o_ (Figure 5). The relaxed length decreases by a large amount in sulfate or PEG and increases by a small amount in perchlorate. *ΔG* is essentially unchanged by any of the co-solvents while the magnitudes of *TΔS, ΔH* and *ΔC*_p_ are all significantly and progressively decreased by large amounts in sulfate and PEG but almost unchanged in 0.3 m perchlorate. In 30% PEG, *Δ*H and *ΔC*_p_ approach zero, *ΔG ∼ - TΔS* (Figure S2), and the ordered water fraction in the NMR experiment is greatly reduced.

## Conclusions

At physiological temperature, recoil in elastin is characterized by several unusual properties: (i) the internal energy increases, (ii) the heat capacity decreases, (iii) the entropic driving force is large compared to the net driving force because spontaneous recoil is highly endothermic and (iv) stretching elastin orders water. Properties (ii), (iii) and (iv) are characteristic of a spontaneous process driven by the hydrophobic effect (29, 30) and all of these properties are different or absent in rubber where the driving force for recoil is the increase in the polymer’s configurational entropy (21, 49). We conclude that at physiological temperatures and in the absence of co-solvents, the predominant driving force for recoil in elastin is the hydrophobic effect.

Natural and artificial configurational entropy springs such as rubber suffer from strain crystallization while experiencing repeated strain cycles, undergoing microscopic phase transitions that result in zones of ordered material with low compliance (50). Hydrophobic effect-driven recoil may be the origin of elastin’s unusual resilience in that it offers a pathway to avoid these phase transitions by retaining disorder even while extended, enabling elastin to be less prone to hysteresis than rubber because water reorientation is more facile than chain motion in a condensed polymer.

## Materials and Methods

### Thermodynamic properties from the temperature dependence of Elastin’s mechanical properties

At constant temperature and pressure, elastin immersed in water is an open system in equilibrium with the surrounding solvent and the relevant thermodynamic expression is the Gibbs free energy (29),

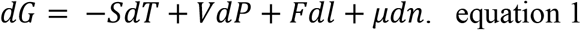

In equation 1, *F* is force, *l* is the sample length and, respectively, *n* and *µ* are the number of moles and chemical potential of water in the elastin matrix. The chemical potential term, *µdn*, accounts for a change in hydration of the elastin matrix when it is stretched or the temperature is changed. Other variables have their usual meaning. It is shown in Figure 3c, and the related discussion, that when elastin is at osmotic equilibrium (*eq*) with the surrounding water, the changes in elastin’s volume and moles of water are related by, 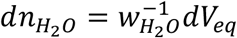, where 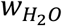 is the partial molar volume of water in the elastin matrix. Since *V*_*eq*_ depends only on temperature and length, *dV*_*eq*_ can be expanded in *dT* and *dl*,

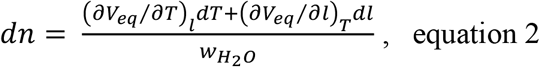

Substituting equation 2 into equation 1 gives a free energy expression subject to the constraint that hydrated elastin is at equilibrium with the water in which it is immersed.

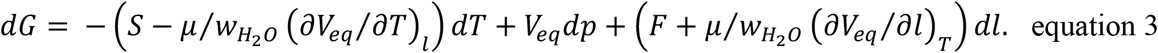

From equation 3, the following Maxwell relation (51) is obtained:

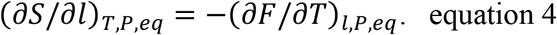

Terms containing *V*_*eq*_ cancel because the order of partial differentiation does not matter. After equation 4 is integrated with respect to *l*, the relation for the entropy change is,

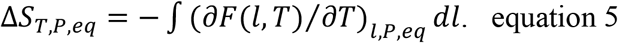

Because the volume change with stretch is small, Figure 3a and (*∂V*_*eq*_*/∂l*)_*T*_ < 4 mm^2^, Δ*G*_*T,P,eq*_ is the integral of the stretch force with respect to *l*,

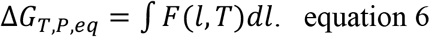

Previously, Flory separated the total strain (*f=*F/a and a = the cross-sectional area) into the sum, respectively, of enthalpic (*f*_*H*_) and entropic (*f*_*S*_) components, (13)

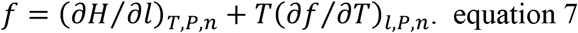

It is shown, in the supplemental information, that equation 7 for a system in equilibrium with the surrounding solvent can be derived directly from equations 4 and 6 and the definition of the Gibbs free energy.

To experimentally determine *F(l,T*), the sample length, *l*, was measured from 3°C to 55°C with forces from 0.09 N to 0.72 N at each temperature. The length was found to accurately fit a virial equation of state that is 1^st^-order in 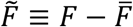 and 3^rd^-order in 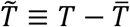.

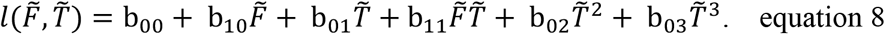

The polynomial, equation 8, is an expansion about the averages of the experimental forces and temperatures. The coefficient b_*1*2_ was found to be zero within experimental error and not included in equation 8. With the definitions of the length at zero force,

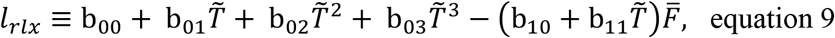

and the spring constant,

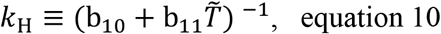

equation 8 can be solved for *F* and written in the form of Hooke’s law,

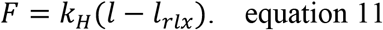

Note that *k*_H_ and *l*_*rlx*_ are temperature-dependent functions of the virial coefficients and are related to the Young’s modulus by *E* = *k*_*H*_*l*_*rlx*_*/a*. They provide greater physical insight than the Young’s modulus alone and all thermodynamic properties can be calculated from them. The free energy (non-*PΔV* work) is obtained by inserting equation 11 into equation 6 and integrating,

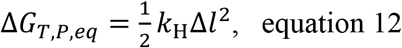

and Δ*l* ≡ *l* − *l*_r*lx*_. Differentiating equation 11 with respect to temperature, we obtain,

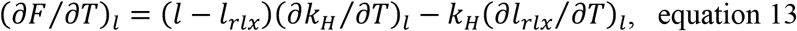

which is inserted into equation 5 and integrated to obtain the entropy change,

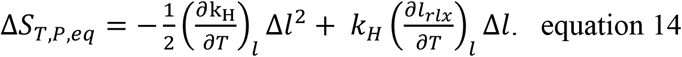

In turn, Δ*H*_*T,P,eq*_, is obtained using the general expression Δ*H* = Δ*G* + *T*Δ*S* and the heat capacity, Δ*C*_*p*_ = (*∂*Δ*H*_*T,P,eq*_/*∂T*)_*P*_, is

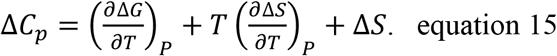

The partial derivatives in equation 15 are obtained from equations 12 and 14 yielding,

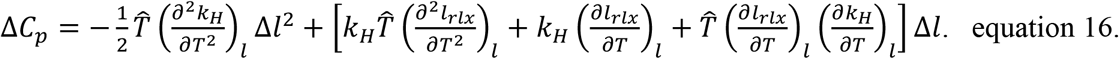

Analytical expressions for the partial derivatives in equations 14 and 16 are readily obtained from the definitions of *l*_*rlx*_ *and k*_*H*_, equations 9 and 10.

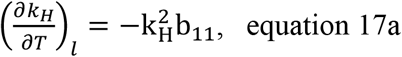

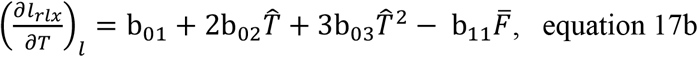

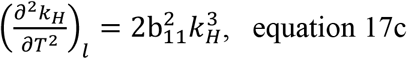

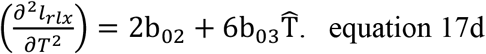

### Elastin sample preparation

A 0.24 g dried elastin fiber (∼ 77 mm × 1.6 mm × 6.4 mm) was cut from purified, bovine neck ligament. The purification procedure,(32) avoids harsh conditions used in older procedures, removes microfibrils, collagen and preserves fibrillar structure of native elastin (Figure 1a). A single sample was used for all of the thermomechanical data reported in Table I and Figures 1-4. After each co-solvent study, the sample was back exchanged into water and a length versus temperature curve (Figure S6) confirmed that the co-solvent had been removed and the sample was unchanged. The sample was glued (Elmer’s E616 Super Glue) to 1/4” diameter rods at both ends for attachment to the stretcher apparatus. A separate and larger sample was used to determine the temperature dependence of elastin’s mass (Figure 6b). With the larger sample (dry mass of 2.029g), the contribution to the total mass from water on the surface of the sample was minimized. After removal from the temperature-controlled water bath, samples were quickly blotted on a chem-wipe before weighing. Additional water loss that occurs on sample warming was minimized by performing the experiments in the cold room.

### Stretcher Instrument

The sample compartment of the apparatus (Figure S4) used to determine *F(l*,T) and *V(l*,T) was machined from delrin (sides and base) and plexiglass windows (front and back) through which the fiber was photographed with a 36-megapixel digital camera (Sony A7r). The short rod glued to the lower end of the elastin sample was fixed to the base and the top rod was attached to a string that was passed over a low friction pulley (Super pulley, ME9450a, Pasco, Roseville, CA) to an adjustable mass that was converted to force using the acceleration due to gravity, 9.8 m/s^2^. The camera and tank were rigidly fixed to a table, from which the fiber length was measured as a function of applied force and temperature with an accuracy of 0.1mm to 0.2mm using ImageJ software (https://imagej.nih.gov/ij/). The digital image was calibrated against a 200 mm ruler placed next to the sample. The temperature controller (blue box in Figure S4c) regulated to within 0.1°C of the set point with a PID controller and the temperature dependence was measured with a platinum resistance thermometer placed in the sample compartment. To avoid thermal gradients, solvent was continuously circulated from the top to the bottom of the tank by a miniature pump.

### Volume as a function of strain and temperature

The sample volume was determined directly from the digital image. Combined with the length measurement, the sample width along the length of the elastin sample was also measured and the sample thickness, the smallest dimension, was calculated from the width measurement with the assumption that the thickness to the width ratio was the same as in the dry sample.

### Thermo-mechanical data collection, analysis and standard error determinations

Length versus force and temperature data were collected as separate *l(F)* runs at 9 forces from 0.09N to 0.72N and *l*(*T*) runs collected at 1°C intervals from 2°C to 55°C. The data were combined into a single table with a typical size of 300 *l*(*F,T*) values from which thermo-mechanical properties were calculated using the two Matlab scripts listed in the Supplemental data. With the first script, the *F(l,T)* data were least-squares fit to equation 8 to determine the virial coefficients, b_ij_, from which all thermo-mechanical properties were calculated in the second script using equations 9-17.

Standard errors were determined by propagating the effects of random errors in (a) the experimental lengths, *l*, or (b) the virial coefficients, b_ij_, into the thermo-mechanical properties (33). Errors in *l* were simulated by adding a random error to each experimental value of *l* sampled from a normal distribution with a standard deviation equal to the accuracy of the measurements in our experimental apparatus, 0.25mm. Multiple data sets simulated in this way were each fit to equation 8, from which a distribution of values was calculated for each thermo-mechanical properties as a function of strain and temperature using equations 9-17.

Alternatively, random errors were added to the coefficients, b_ij_, using the standard errors, σ_ij_, from the fit of the experimental data to equation 8, Table S1, and distributions of properties were calculated as in (a). Standard deviations of the distributions converged with 1000 simulations and are used as standard errors for all properties. To avoid over interpreting the results, the two-fold larger standard errors obtained with method (b) have been used throughout and are shown as error bars at the regular intervals in Figures 2, 4, S1, S2 and S3. Standard errors in Table 1 were calculated in the same way.

### Strain induced changes in solvent ordering studied by double quantum ^2^H NMR

We have observed the effect of stretch on ordering of ^2^H_2_O in the elastin matrix. Insofar as molecular reorientation of ^2^H_2_O is anisotropic, as is the case for ordered water, ^2^H_2_O has a non-zero ^2^H quadrupole coupling and the “forbidden” double quantum (2Q) NMR transition can be observed using the methods of indirect detection. (28) Because reorientation in bulk water is isotropic, the quadrupole coupling is averaged to zero, the 2Q transition is absent and its NMR signal can be removed by 2Q filtration and only the 2Q transition is observed (26, 27). To determine the effect of strain on the fraction of ordered water in the sample, f_o_, we observed the buildup of the 2Q NMR signal from ordered water, *I*_*2q*_, as a function of the 2Q preparation time, t_1_. The NMR signal is proportional to f_o_ and is well-described by the relation, (26)

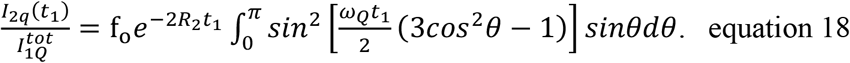

The ordered fraction, f_o_, and the residual quadrupole coupling, *v*_*Q*_ = *ω*_*Q*_/2*π*, were determined by a two parameter least-squares fit of *I*_2*q*_(t_1_) to equation 18 with 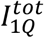, the 1Q signal from all of the ^2^H_2_O in the sample, and *R*_2_, the transverse relaxation time, constrained to the values separately determined in Hahn echo experiments with and without 2Q filtration (Figure S5) (26).

2Q filtered ^2^H spectra were collected on a homebuilt, 11.6 T instrument with a solenoidal coil probe (52) (90° pulse width = 4.2 µs) and the previously described pulse sequence and phase cycle (26). ^2^H spectra (2k complex points) were obtained with a 4 or 5 kHz offset and 128 equally spaced 2Q preparation times, t_1_. Two-dimensional spectra were Fourier transformed in the observed dimension, t_2_, and peak heights of the 2Q filtered signals are displayed (Figure 5) as a function of the 2Q preparation time, t_1_. Suppression of the 1Q signal from bulk water to a level below the noise was confirmed in two ways. All NMR signals in Figure 5 at zero 2Q preparation times (t_1_ = 0) are, within the S/N ratio of the experiments, also zero and 1Q bulk water signals would have been observed only if 2Q filtration was insufficient, as was not the case. In the 2D spectrum, Figure S6, the 1Q (bulk water) and 2Q (ordered water) signals are separated in the F_1_ dimension (26). With a 4 kHz offset, the 2Q peak at F_1_ = 8 kHz from ordered water is large (S/N = 47:1) and any 1Q peak from bulk water at F_1_ = 4 kHz is reduced to below the noise level.

Suppression of the 1Q signal from bulk water to a level below the noise was confirmed in two ways. In the 2D spectrum, Figure S7, the 1Q (bulk water) and 2Q (ordered water) signals are separated in the F_1_ dimension and the phase cycle for eliminating the bulk water (1Q) signal is applied (26). With a 4 kHz offset, the ordered water peak in the indirect dimension at F_1_ = 8 kHz is large (S/N = 47:1) and the bulk water peak at the 1Q frequency, F_1_ = 4 kHz, is reduced to below the noise level. Furthermore, in each of the graphs shown in Figure 5, the 2Q signal at short 2Q preparation times is below the noise level and this is only possible if the much larger bulk water signal is adequately suppressed by the phase cycle.

### NMR sample preparation

Both ends of a purified elastin fiber (∼20 mg and ∼1mm × ∼12 mm) were superglued to ∼12 mm lengths of 1/16” diameter plastic rod, equilibrated overnight in the desired aqueous solution and inserted into a 3 mm NMR tube (length ∼30 mm) after removal of excess solvent by blotting on paper. With the elastin sample centered in the NMR tube, one rod was sealed and held stationary to the tube with glue and the other rod was sealed and held in place with parafilm which could be removed so that the fiber could be stretched to a different length and resealed. Fiber lengths were measured with a digital caliper.

### Materials

Elastin samples were purified from fresh bovine nuchae membranes using a method that preserves the natural fibrous structure and removes other proteins (Figure 1a) (32). Elastin was washed in aqueous sodium chloride for three days to remove soluble proteins followed by ethanol, chloroform, methanol and acetone extraction to remove lipids. Because elastin has no methionine residues and the ε−amino groups of lysyl residues are oxidized for cross-linking, remaining proteins were removed by washing after proteolysis with cyanogen bromide and trypsin. This material has nearly the same modulus at 24°C as was measured in unpurified, single elastin fibers (53). Dry fibers were stored at – 80 °C. Sample solutions were prepared from deionized water with reagent grade solutes (Na_2_SO_4_, 20 kDa PEG, and NaClO_4_). The experiments in deuterated solvent used 99% ^2^H_2_O (Cambridge Isotopes, Andover, MA).

## Supporting information

Supplemental data for "Elastin Recoil is Driven by the Hydrophobic Effect'

## Author Contributions

N.M.J. and R.J.W. designed and performed research; N.M.J, R.L.K and R.J.W. wrote the manuscript.

## Notes

The authors declare no competing financial interests.

## Acknowledgements

The authors gratefully acknowledge support from the NSF (DMR-1410678) to R.J.W and R.L.K and help in fitting the NMR data from Dr. Abed Jamhawi.

## Abbreviations

NMR: nuclear magnetic resonance
2Q: double quantum
PEG: polyethylene glycol

## Data availability statement

All data used in Figures 1-5 and Table 1 and the supplemental information is available at https://academicworks.cuny.edu/cc_pubs/985/. All thermomechanical data has been distilled into the virial coefficients listed in SI Table 1 from which properties in Figures 2, 3c and 4 were calculated as described in the materials and methods sections.

## Code availability statement

The custom Matlab scripts used to calculate the virial coefficients from the raw thermodynamic data and the thermomechanical properties and their associated confidence intervals from the virial coefficients are in the supplemental information.

