## Supplemental data for "Elastin Recoil is Driven by the Hydrophobic Effect' for "Elastin Recoil is Driven by the Hydrophobic Effect"

### Supplemental Information

**S1:** The virial coefficients for the equation of state, equation 6, for all studied solutions.

| Solution | $\overline{F}$ (N) | $\overline{T}$(K) | $b_{00}$ ($\times10^{-3}$) | $b_{10}$ ($\times10^{-2}$) | $b_{01}$ ($\times10^{-4}$) | $b_{11}$ ($\times10^{-5}$) | $b_{02}$ ($\times10^{-6}$) | $b_{03}$ ($\times10^{-8}$) |
|---|---|---|---|---|---|---|---|---|
| $^1H_2O$ | 0.38 | 301.31 | 77.55 ± .04 | 1.82 ± .01 | -2.4 ± .04 | -8.65 ± .83 | 7.17 ± .13 | -6.91 ± .79 |
| $^2H_2O$ | 0.38 | 302.08 | 76.73 ± .05 | 1.82 ± .02 | -1.57 ± .06 | -8.05 ± 1.13 | 6.66 ± .21 | -16.47 ± 1.54 |
| 0.1 m $Na_2SO_4$ | 0.38 | 301.68 | 75.79 ± .04 | 1.81 ± .01 | -1.70 ± .04 | -8.04 ± .73 | 6.35 ± .12 | -11.78 ± .86 |
| 0.3 m $Na_2SO_4$ | 0.37 | 300.58 | 73.94 ± .03 | 1.77 ± .01 | -1.23 ± .03 | -7.67 ± .60 | 4.04 ± .01 | -6.32 ± .66 |
| 15% 20 kDa PEG | 0.38 | 299.75 | 76.37 ± .07 | 1.73 ± .02 | -1.53 ± .07 | -6.49 ± 1.43 | 4.28 ± .25 | -5.45 ± 1.61 |
| 30% 20 kDa PEG | 0.38 | 300.07 | 73.86 ± .04 | 1.66 ± .01 | -0.33 ± .04 | -2.50 ± .88 | 1.00 ± .15 | -0.52 ± 1.00 |
| 0.3 m $NaClO_4$ | 0.38 | 301.06 | 80.20 ± .02 | 1.85 ± .01 | -2.35 ± .03 | -6.10 ± .55 | 5.16 ± .01 | -4.85 ± .64 |
| 1.0 m $NaClO_4$ | 0.38 | 301.07 | 79.58 ± .04 | 1.76 ± .01 | -1.47 ± .04 | -6.33 ± .85 | 3.29 ± .15 | -4.79 ± 1.03 |

*Complete Thermomechanical Data*

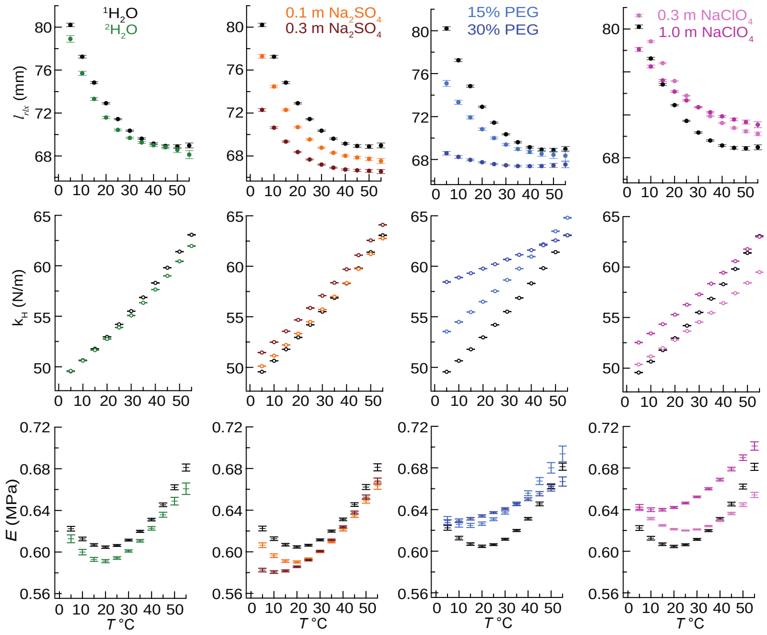


**S2:** Temperature dependence of elastin's mechanical parameters (rows) versus strain in different solvents (columns): $^1H_2O$ (black) and $^2H_2O$ (green) are in column **a**, $Na_2SO_4$ (brown) in column **b**, PEG (blue) in column **c** and $NaClO_4$ (magenta) in column **d**. The co-solvents are dissolved in $^1H_2O$ and darker shading indicates the higher concentration.

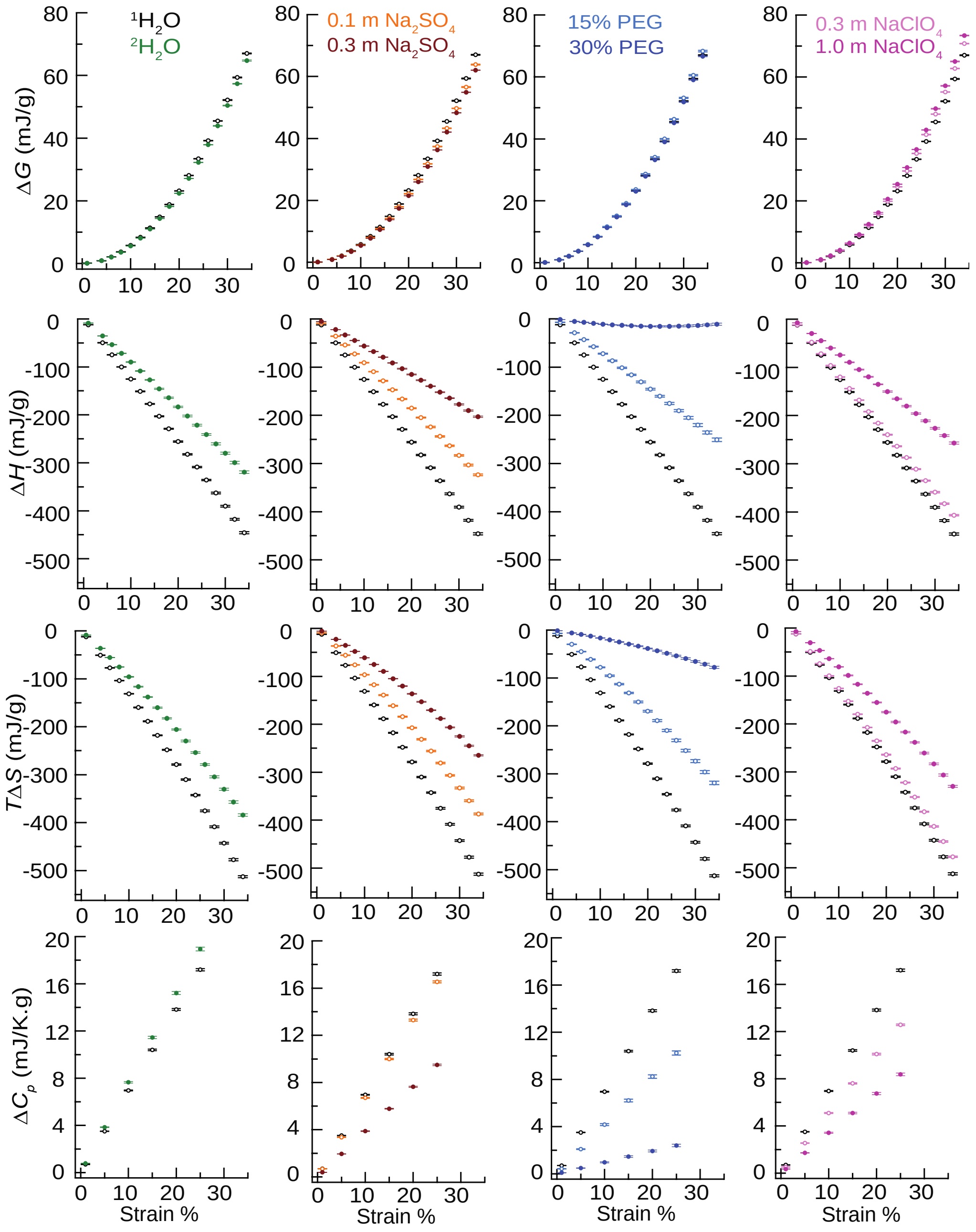


**S3:** Strain dependence of Elastin's thermodynamic parameters (rows) versus strain at 25°C in different solvents (columns): $^1H_2O$ (**o**). $^2H_2O$ (o), $Na_2SO_4$ (o), PEG (o) and $NaClO_4$ (o). The co-solvents are dissolved in $^1H_2O$ and darker shading indicates the higher concentration.

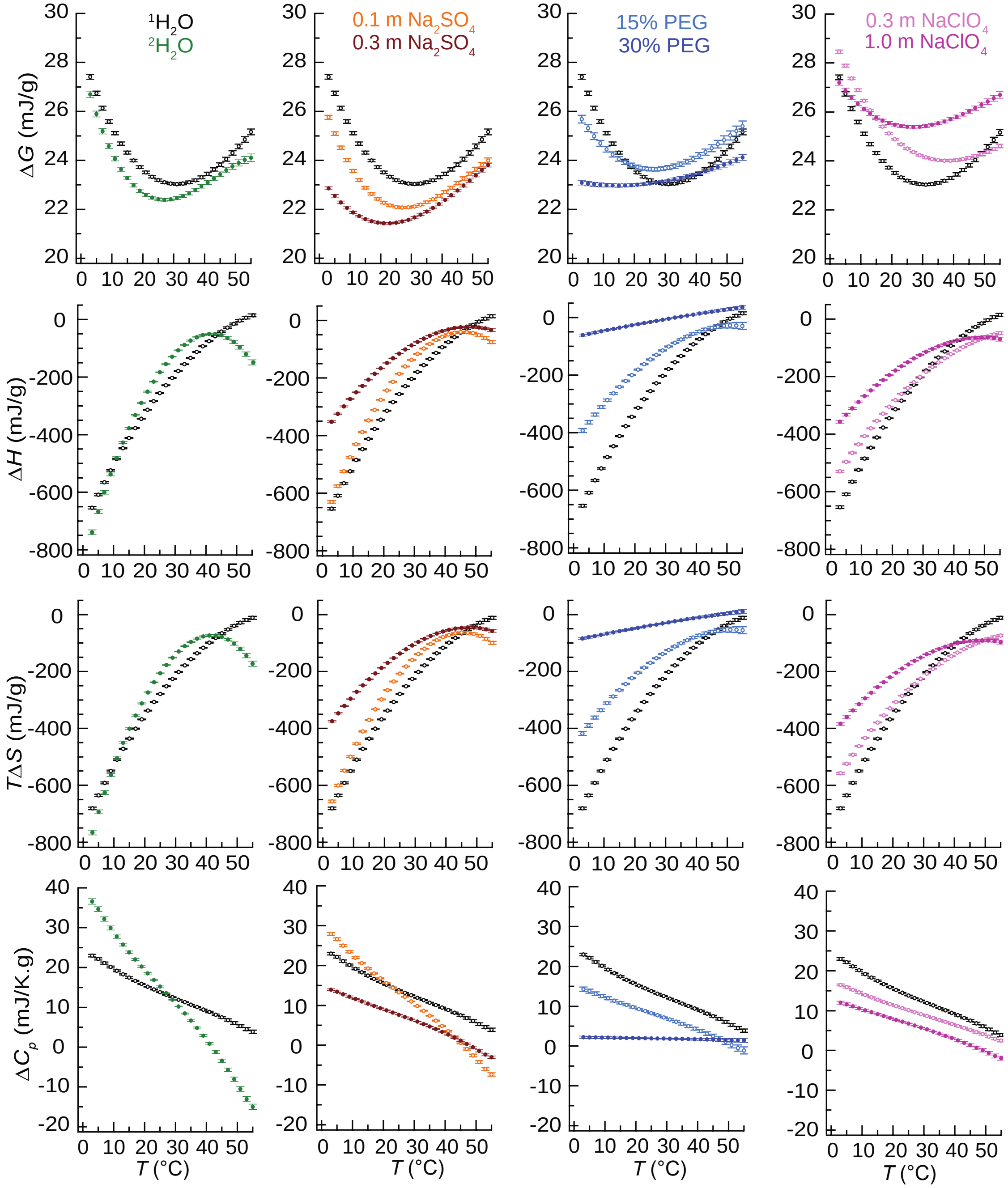


**S4:** Temperature dependence of elastin's thermodynamic parameters at 20% strain in different solvents (columns): $^1H_2O$ (**o**), $^2H_2O$ (o), $Na_2SO_4$ (o), PEG (o) and $NaClO_4$ (o). The co-solvents are dissolved in $^1H_2O$ and darker shading indicates the higher concentration.

**S5**: Equation 7 from the main text, proposed by Flory without a derivation, can be obtained from the definition of the Gibbs free energy, $G \equiv H - TS$ and equations 4 and 6 which were derived in the main text. The partial derivative of $G$ with respect to $l$ at constant $T$, $P$ and at osmotic equilibrium is,

$$\left(\frac{\partial G}{\partial l}\right)_{T,P,eq} = \left(\frac{\partial H}{\partial l}\right)_{T,P,eq} - T\left(\frac{\partial S}{\partial l}\right)_{T,P,eq}. \quad \text{SI equation 1}$$

From the main text, equation 4 and equation 6 before integration are,

$$(\partial S/\partial l)_{T,P,eq} = -(\partial F/\partial T)_{l,P,eq} \text{ and } (\partial G/\partial l)_{T,P,eq} = F(l,T). \quad \text{SI equations 2}$$

By inserting SI equations 2 into SI equation 1,

$$\left(\frac{\partial F}{\partial l}\right)_{T,P,eq} = \left(\frac{\partial H}{\partial l}\right)_{T,P,eq} + T\left(\frac{\partial F}{\partial l}\right)_{T,P,eq}, \quad \text{SI equation 3}$$

and Flory's equation for the uniaxial strain, $f$, is obtained after dividing both sides of SI equation 3 by the cross-sectional area,

$$\left(\frac{\partial f}{\partial l}\right)_{T,P,eq} = \left(\frac{\partial H}{\partial l}\right)_{T,P,eq} + T\left(\frac{\partial f}{\partial l}\right)_{T,P,eq}. \quad \text{equation 7}$$

Note that in equation 7, $H$ is normalized by the cross-sectional area.

*Stretcher apparatus*

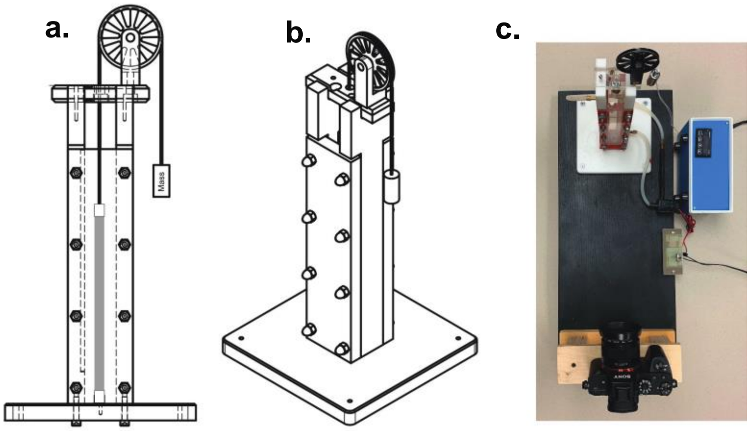


**S6**: Stretcher Instrument sample compartment (a) front view and (b) 3-D perspective. The elastin sample is shaded gray. (c) Top-down photograph of the instrument with the camera (bottom), sample compartment (top) and T-controller (top right).

*Relaxation data used in the analysis of the 2Q NMR data.*

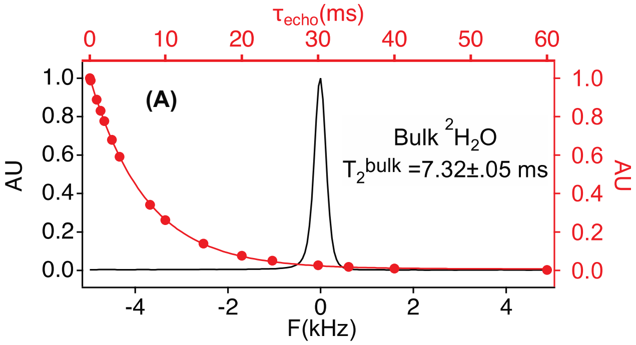


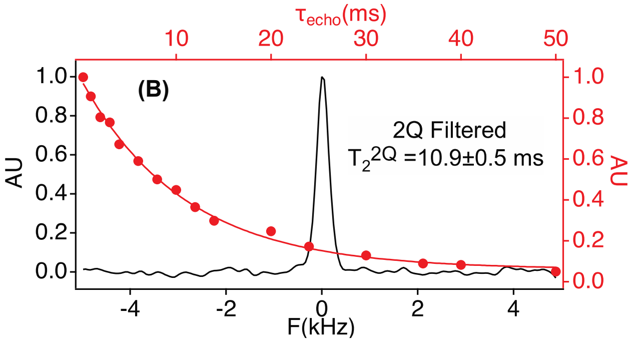


**S7:** Determinations of the $^2H_2O$ relaxation rates, $T_2$ (A) and $T_{2Q}$(B), respectively, using a Hahn echo (A) and a Hahn echo with 2Q filtration (B). Elastin was hydrated in $^2H_2O$. The spectra shown were those obtained with an echo delay of 4 ms and are representative of the other relaxation measurements summarized in S8. Spectra in (A) and (B) were obtained with 8 and 128 transients, respectively. The lower S/N, ~ 20, of the spectrum in (B) compared to that in (A), S/N ~ 100, is due to signal loss during the $\mathbf{1Q} \leftrightarrow \mathbf{2Q}$ transfers in the $\boldsymbol{T_2^{2Q}}$ experiment (1).

**S8**: $^2H_2O$ NMR $T_2$ relaxation data obtained from elastin at different stretch values and in $^2H_2O$, 0.1 m $Na_2SO_4$ and 10% PEG. $T_2^{bulk}$ and $T_2^{2Q}$ are $T_2$ values measured, respectively, with and without 2Q filtration

| **Solution** | **% stretch** | $T_2^{bulk}$ (ms) | $T_2^{2Q}$ (ms) |
|---|---|---|---|
| $^2H_2O$ | 0% | 13.1 ± 0.3 | 12.4 ± 0.5 |
| " | 15 % | 12.2 ± 0.2 | 10.4 ± 0.1 |
| " | 33 % | 9.7 ± 0.2 | 11.3 ± 0.1 |
| 0.1 m $Na_2SO_4$ | 0 % | 10.6 ± 0.2 | 9.4 ± 0.2 |
| " | 14 % | 10.6 ± 0.3 | 9.7 ± 0.2 |
| " | 28% | 10.5 ± 0.3 | 9.2 ± 0.1 |
| 10% PEG | 0 % | 9.3 ± 0.3 | 8.4 ± 0.3 |
| " | 15 % | 8.9 ± 0.2 | 7.3 ± 0.1 |
| " | 26 % | 7.7 ± 0.1 | 6.6 ± 0.1 |

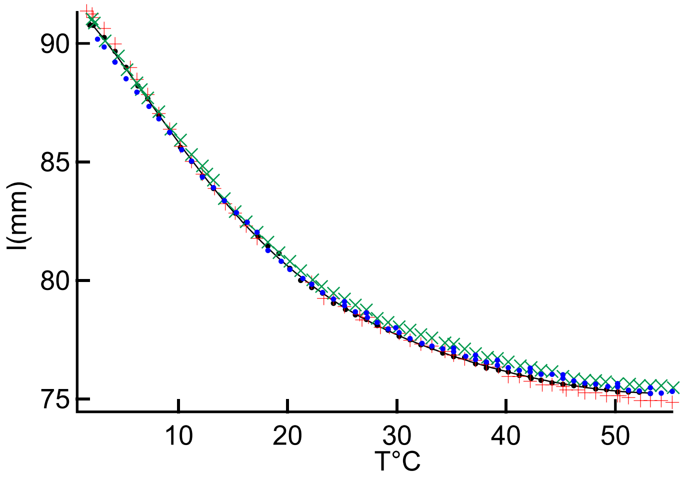


**S9**: Elastin sample lengths vs temperature at 0.40 N force data obtained after the elastin sample in the indicated co-solvent was back exchanged into water. $^1H_2O$ (black), $Na_2SO_4$ (red), PEG (green), $^2H_2O$ (blue)

*Raw data files available at the indicated URL*

**S10**: URL for access to raw thermodynamic data, 2Q filtered $^2H$ NMR data and the volume data.

https://academicworks.cuny.edu/cc_pubs/985/

**References, SI**


1. T. V. Krivokhizhina, R. Wittebort, 2Q NMR of $^2H_2O$ ordering at solid interfaces. *Journal of Magnetic Resonance* **243**, 33-39 (2014).
